# Unexpected Molecular Mechanism of Orc6-Based Meier-Gorlin Syndrome: Insights from a Humanized Drosophila Model

**DOI:** 10.1101/2025.03.31.646466

**Authors:** Maxim Balasov, Katarina Akhmetova, Igor Chesnokov

## Abstract

Meier-Gorlin syndrome (MGS) is a rare autosomal recessive disorder characterized by microtia, primordial dwarfism, and skeletal abnormalities. Patients with MGS often carry mutations in genes encoding the subunits of the Origin Recognition Complex (ORC), components of the pre-replicative complex and replication machinery. ORC6, an essential ORC subunit, plays a critical role in both DNA replication and cytokinesis. Approximately 30% of reported ORC6-related MGS cases exhibit compound heterozygosity for the ORC6 variants c.2T>C (p.Met1Thr) and c.449+5G>A. The c.2T>C mutation disrupts the start ATG codon by changing it to ACG, potentially initiating translation at an alternative downstream in frame Methionine (Met20), while c.449+5G>A results in in-frame exon skipping. Both mutations are predicted to produce significantly truncated ORC6 proteins with impaired functionality. In this study, using humanized ORC6 based *Drosophila* model, we demonstrated that these truncated proteins fail to rescue *orc6* deletion. Instead, our findings reveal that the strong Kozak sequence, naturally present in human ORC6 mRNA, promotes translation from a non-canonical ACG codon. Rescued flies demonstrated a phenotype that we observed earlier for other MGS mutants in *Drosophila*. These results provide compelling evidence that MGS patients with c.2T>C/c.449+5G>A mutation rely on full size ORC6 protein initiated from a non-canonical ACG start codon.

## Introduction

Meier-Gorlin syndrome (MGS) is a rare form of microcephalic primordial dwarfism. It is also known as ear, patella, short stature syndrome and/or microtia, absent patella, and micrognathia syndrome, traits highlighting the core clinical phenotypes ^1-4^. Generally, this disorder is inherited in an autosomal recessive manner. Genes linked to MGS are involved in processes required for cell cycle progression such as cell cycle checkpoint, DNA replication, and DNA repair ^5^. The development of high throughput sequencing methods, along with an increase in patient cohorts, contributed significantly to both the genetic and phenotypic spectrums of MGS. However, the molecular mechanisms responsible for the manifestation of specific mutations remain to be elucidated for most of the genes.

Variants in a number of DNA replication genes have been associated with MGS including genes encoding Origin Recognition Complex (ORC) and MCM helicase subunits, CDT1, CDC6, and other components of the pre-replicative complex and replication machinery ^6^ (and references therein), suggesting that the clinical phenotypes are caused by defects in DNA replication. DNA replication is fundamentally important for tissue development, growth and homeostasis, and impairments of the DNA replication machinery have catastrophic consequences for genome stability and cell division. ORC6 is a component of ORC important for the initiation of DNA replication in all species ^7-13^. Metazoan ORC6 protein consists of two functional domains: a larger N-terminal domain important for DNA binding, and a smaller C-terminal domain important for protein-protein interactions ^9,11,14-17^. In our previous studies we established fly models of MGS and have shown that the Y225S MGS mutation in Orc6 C-terminus ^2,3^ impaired the association of Orc6 with the core ORC1-5 complex ^16,18^; another MGS mutation in the protein N-terminus, K23E, ^19^ directly affected DNA binding ^20^.

Approximately one-third of reported individuals with causal variants in *ORC6* have compound heterozygosity in the *ORC6* gene: c.2T>C (p.Met1Thr) and c.449+5G>A (splice site mutation) ^6^. These MGS patients have relatively mild phenotypes similar to the majority of *ORC6*-related MGS cases. Mutation c.2T>C disrupts the start ATG codon by changing it to ACG, and is thought to cause initiation at a downstream in frame Methionine (Met20), leading to the production of a truncated protein missing first 19 amino acids of the N-terminal domain ^6^. The c.449+5G>A mutation has been shown to cause in-frame exon 4 skipping *in vitro*, which presumably results in the deletion of 30 amino acids from the middle of the N-terminal domain ^27^. Exon skipping is not the only possible outcome of the 5G>A mutations of a splice donor site. It has been shown that the mutation of this type might lead to the reduced splicing efficiency ^21^ suggesting a possibility that c.449+5G>A may lead to the overall decreased expression level of the ORC6 protein product.

Our earlier structural studies and molecular genetic analyses ^15,17^, along with functional screens using CRISPR-Cas9 techniques ^22^, indicate that the N-terminal domain of ORC6 is essential for maintaining ORC complex functionality. Large deletions in this domain are expected to produce non-functional proteins. However, the relatively mild phenotype observed in patients with the MGS mutations c.2T>C and c.449+5G>A suggests that the resulting truncated proteins should retain some residual functionality ^6^.

In this study, we used humanized *Drosophila* models to test various possible outcomes of c.2T>C and c.449+5G>A variants *in vivo*. Our findings indicate that truncated proteins predicted to result from both variants are nonfunctional. We also tested the effect of c.449+5G>A on splicing efficiency and found that it completely abolishes splicing, with no mRNA or protein product detected. Instead, we discovered that the human *ORC6* gene naturally contains a classical Kozak sequence ^23^ which facilitates the translation of a full-length ORC6 protein from a non-AUG start codon ACG (c.2T>C mutation) resulting in transgenic flies which show phenotypes previously reported for other MGS mutations ^18,20^. Using genetic tools available for *Drosophila*, we increased the expression of the c.2T>C mutant transgene, which restored the MGS phenotype in flies to the wild type. These results strongly support the hypothesis that the mild MGS phenotype observed in patients with biallelic missense variants c.2T>C and c.449+5G>A is due to a reduced amount of the full-length ORC6 protein.

## Materials and Methods

### Mutagenesis and cloning

Human-*Drosophila* hybrid Orc6 contains first 206 amino acids of human ORC6 and last 52 amino acids of *Drosophila* Orc6, and was described in our previous work ^20^. Deletions in hybrid *orc6* were generated by PCR technique and cloned in frame with GFP tag at the N-terminus. Intron 4 was recovered from Human Genomic DNA (#11691112001, Roche) and inserted into human-*Drosophila* hybrid at its native position. Intronic mutation c.449+5G>A was created by site-directed mutagenesis. For mimicking the DNA sequence context of the Meier-Gorlin mutation c.2T>C, the nine upstream nucleotides from human mRNA (https://www.ncbi.nlm.nih.gov/nuccore/AF139658.1) were added to human-*Drosophila* hybrid carrying mutation c.2T>C. Intronic and mimic transgenes were fused with FLAG peptide at the C-terminus. All hybrid genes were cloned into the modified *pUAST* vector containing both inducible *UAS* promoter and *Drosophila orc6* native promoter, verified by sequencing and injected into *Drosophila w*^*1118*^ embryos (Model System Injections, Raleigh, NC). Individual transgenic strains were set up.

### Fly stocks and rescue experiments

Fly stocks containing endogenous *orc6*^*35*^ deletion ^13^ and N-terminus GFP fused transposon:

| Genotype | Genetic modification | Abbreviations used in paper |
| --- | --- | --- |
| *orc6<sup>35</sup>/Cy; HDWT* | Hybrid protein containing first 206 amino acids of human ORC6 and last 52 amino acids of *Drosophila Orc6* <sup>20</sup> | HDWT |
| *orc6<sup>35</sup>/Cy; HDM20* | Deletion of first 19 amino acids in human-*Drosophila* hybrid | HDM20-1<br>HDM20-2<br>Two independent transgenic stocks |
| *orc6<sup>35</sup>/Cy; HDdel* | Deletion of the 30 amino acids between Serine 120 and Lysine 150, Serine is also mutated to Arginine. | HDdel-1<br>HDdel-2<br>Two independent transgenic stocks |

Fly stocks containing endogenous *orc6*^*35*^ deletion and C-terminus FLAG peptide fused transposon:

| Genotype | Genetic modification | Abbreviation used in paper |
| --- | --- | --- |
| *orc6<sup>35</sup>/Cy; HDIn4* | Wild type intron 4 inserted in native position into human part of hybrid gene | HDIn4-1<br>HDIn4-2<br>Two independent transgenic stocks |
| *orc6<sup>35</sup>/Cy; HDIn4G>A* | Mutant intron 4 (c.449+5G>A) inserted in | HDIn4G>A-1<br>HDIn4G>A-2 |
|  | native position into human part of hybrid gene | Two independent transgenic stocks |
| *orc6<sup>35</sup>/Cy; HD2T>C* | GCCGGCGCC precedes the non-canonical start codon ACG (2T>C mutation). | HD2T>C-1<br>HD2T>C-2<br>Two independent transgenic stocks |

Bloomington Drosophila Stock Center stock #5138 (*y,w*; *P{w*^*+mC*^ *= tubP-GAL4}LL7/TM3,Sb,Ser)* expressing GAL4 ubiquitously under the control of the *alphaTub84B* promoter was used to design *orc6*^*35*^*/Cy*; *Tubulin-GAL4/TM3,Sb* fly stocks for rescue experiments.

In rescue experiments with a native *Drosophila* promoter, progeny from heterozygous *orc6*^*35*^*/Cy*; *HD* was analyzed for the presence of *orc6*^*35*^*/orc6*^*35*^; *HD* adult flies. In rescue experiments with *alphaTub84 (Tubulin)* promoter, females of the genotype *orc6*^*35*^*/Cy*; *HD* were crossed to males *orc6*^*35*^*/Cy*; *Tubulin-GAL4/TM3,Sb*, and resulting progeny were analyzed for the presence of *orc6*^*35*^*/orc6*^*35*^;*HD/Tubulin-GAL4* adults.

### Mitotic chromosome preparation

Preparation of mitotic nuclei was described previously ^24^. Briefly, third instar larval neural ganglia were incubated in 0.075M KCl for 5 min, fixed in methanol with acetic acid (3:1) for 20 min, and then dispersed in a drop of 50% propionic acid on a slide. Then slides were dried and stained with 5% Giemsa’s solution.

### Western blot

Heterozygous flies of the following phenotype were used for western blot: *orc6*^*35*^*/Cy; HD/Tubulin-GAL4*. Five adult flies or four pairs of ovaries were homogenized in 30 μl of the 300 mM NaCl, 0.5% NP-40 PBS buffer. The samples were centrifuged 5 min at 10000 × g and supernatants were used for Western blotting. For detecting GFP tagged hybrids, anti-GFP mouse monoclonal antibody was used (#SC-9996, Santa Cruz Biotechnology Inc). FLAG-tagged hybrids were detected with anti-FLAG mouse monoclonal antibody (#SC-166355, Santa Cruz Biotechnology Inc). Ubiquitously expressed septin Pnut served as a loading control. Anti-Pnut rabbit polyclonal antibodies were purified in the lab ^25^.

### RNA purification and cDNA generation

RNA was isolated from 5 females using ZR Tissue and Insect RNA MicroPrep (#R2030; Zymo Research). DNA was removed using TURBO DNase (#AM2238; Invitrogen) following manufacturer’s recommendations. cDNA was generated from 1 μg of total RNA using ProtoScript II First Strand cDNA Synthesis Kit (#E6560; New England Biolabs). For detecting human-*Drosophila* hybrid *orc6* cDNA we used primers: TATATCTAGAATGGGGTCGGAGCTGATCGGG (human forward) and GTCAAGCTTCTAAGCCTCGAGAAGCTGG (*Drosophila* reverse). Endogenous *orc6* cDNA was detected with primers AGGTCTAGAATGACTACCTTAATAG (*Drosophila* forward) and GTCAAGCTTCTAAGCCTCGAGAAGCTGG (*Drosophila* reverse). 100bp DNA ladder (#D001, GoldBio) was used in Supplementary Figure 1. 1Kb DNA ladder (#N3232, New England BioLab) was used in Figure 4B, D.

### Expression of human-*Drosophila* hybrid transgene in S2 cells

FLAG-tagged human-*Drosophila* hybrid *orc6* transgenes carrying 9 upstream nucleotides with Kozak sequences, **G**CC**G**GC**G**CCACG**G**, or Kozak sequence with mutated purines in positions -3, -6, -9, **C**CC**C**GC**C**CCACG**G**, were cloned into PMT/V5 vector and transfected into S2 cells using *Trans*IT-insect transfection reagent (#MIR 6100, Mirus Bio) following manufacturer’s recommendations. S2 cells were cultured at 27°C in Shields and Sang M3 medium (#S8398, Sigma-Aldrich) supplemented with 5% fetal bovine serum. After 24h methallothionein promoter was activated by 0.5mM CuSO_4_. On the next day cells were collected and used for western blot and cDNA preparation.

## Results

### Design of the MGS mutations in *Drosophila*

The metazoan ORC6 protein consists of two functional domains, a larger N-terminal domain that has a homology with TFIIB transcription factor and is important for DNA binding ^9,11,15^, and a shorter C-terminal domain that interacts with the core ORC through the Orc3 subunit ^16^. In our previous work we demonstrated that a construct containing human N-terminal domain of ORC6 (∼80% of the protein length) and *Drosophila* C terminus, Human-*Drosophila* (HD) hybrid, rescued lethality associated with *orc6* deletion in *Drosophila*, and resulting flies’ phenotypes were undistinguishable from the wild type ^24^. We successfully used this humanized *Drosophila* model to define molecular consequences of the Meier-Gorlin mutation K23E ^20^ indicating that mutations in the human part of the Human-*Drosophila* hybrid can be explored using *Drosophila* system.

In this study we investigated the molecular basis of MGS phenotypes caused by the compound heterozygosity c.2T>C(p.Met1Thr)/c.449+5G>A ^6^. The mutation c.2T>C in start codon has been thought to cause the initiation of translation at the next downstream Methionine at position 20 resulting in ORC6 protein truncated at the N-terminus ^6^. The intronic c.449+5G>A variant has been shown *in vitro* to cause an in-frame exon skipping of exon 4, which would be translated into an ORC6 protein lacking 30 amino acids in the DNA binding domain (amino acids 120 to 150) ^26^. An alternative possibility for the c.449+5G>A mutation is that it might reduce splicing efficiency of the wild type ORC6 mRNA ^21^.

We used the humanized Human-*Drosophila* (HD) fly system to create several constructs recapitulating human mutations **(Figure 1)**. The first, HDM20, lacks 19 amino acids of the N-terminal domain as a result of Methionine at position 20 acting as a start codon **(Figure 1A, HDM20)**. The second construct, HDdel, carries deletion of 30 amino acids of the middle domain between Serine 120 and Lysine 150. Serine 120 was also mutated to Arginine to mimic the skipping of exon 4 **(Figure 1A, HDdel)** as described in the clinical case ^26^. The third and fourth constructs were designed to test splicing efficiency *in vivo*. The third construct, HDIn4, was generated by inserting wild-type human intron 4 into the human part of the HD hybrid at the native position (**Figure 1B, HDIn4)**. This transgene served as a control to ensure that *Drosophila* splicing machinery can handle human intron. The fourth construct, HDIn4G>A, was constructed by introducing the splice site mutation c.449+5G>A into the intron 4 of the HDIn4 hybrid **(Figure 1B, HDIn4G>A)**. HD wild type hybrid transgene, HDWT, was used as a control since it compensates for *orc6* deletion and restores wild type phenotype ^24^ (**Figure 1A, HDWT)**. All constructs were injected into *Drosophila* embryos and homozygous fly strains were set up for detailed analyses.

**Figure 1.**
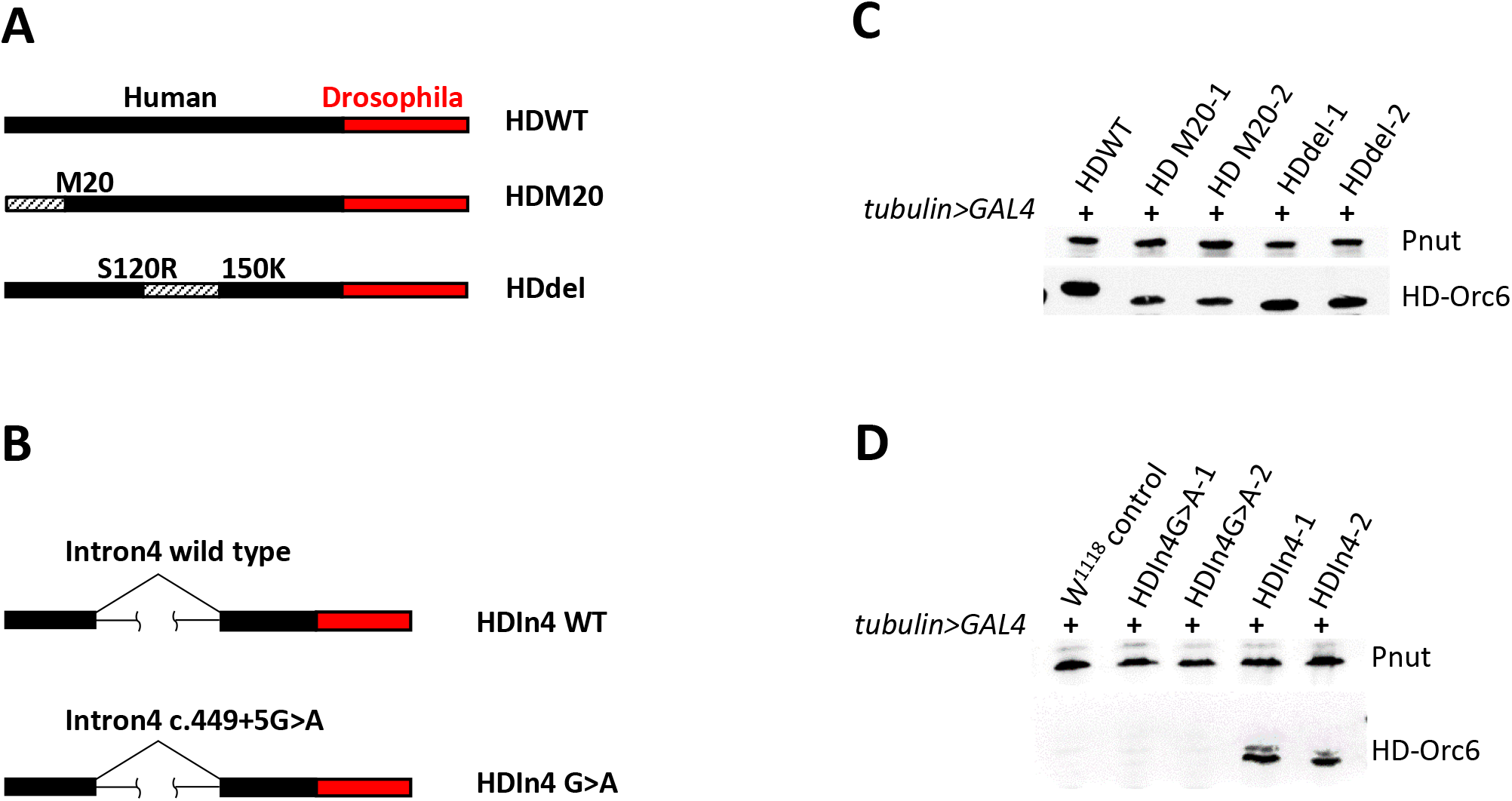
Strategy for the modeling of the Meier-Gorlin syndrome associated with compound heterozygosity in *ORC6* gene (c.2T>C(p.Met1Thr)/c.449+5G>A). **(A)** Mutation c.2T>C causes deletion of the first 19 amino acids as a result of using second Methionine codon at the position 20 (HDM20); c.449+5G>A induces exon skipping and generates deletion of the 30 amino acids between Serine 120 and Lysine 150, Serine is also mutated to Arginine (HDdel). Black bar represents human ORC6 and red bar represents *Drosophila* Orc6. Dashed bar shows deleted part of the protein. **(B)** Wild type intron 4 (HDIn4 WT) or intron 4 containing mutation c.449+5G>A (HDIn4 G>A) was inserted into Human-*Drosophila* hybrid transgene to test splicing efficiency. **(C**,**D)** Western blots of whole flies expressing indicated HD constructs under control of the *tubulin* promoter. Pnut protein was used as a loading control. **(C)** Independent transgenic fly stocks HDM20-1 and HDM20-2 have first 19 amino acids deleted. HDdel-1 and HDdel-2 have missing 30 amino acids of the DNA binding middle domain of ORC6. HD wild type (HDWT) is a full size 258 amino acids Human-*Drosophila* hybrid. **(D)** The HDIn4G>A-1 and HDIn4G>A-2 are two independent fly stocks with mutation in splice site c.449+5G>A. The HDIn4-1 and HDIn4-2 are independent fly stocks bearing transgene with wild type intron 4 (positive control). *w*^*1118*^ represents negative control.

### The analysis of flies carrying mutated Human-*Drosophila* hybrids

Replication defects often result in a loss of chromosome integrity. In our previous studies we showed that cytological investigation of mitotic chromosomes in developing larval brains provides a convenient tool for monitoring the effects of transgenic constructs bearing mutations of interest ^18,20^. The normal *Drosophila melanogaster* karyotype consists of four pairs of chromosomes (**Figure 2A**). For quantification of chromosome integrity, we counted percentages of normal mitoses, mildly/moderately affected and severely affected mitoses (**Figure 2A**). Abnormal mitoses are observed in flies with *orc6* deletion, or flies expressing full length human Orc6 (HsOrc6) on *orc6* deletion background. In contrast, the expression of the Human-*Drosophila* wild type hybrid transgene (HDWT) supports normal mitotic configuration as wild-type *Drosophila* Orc6 (DmWT) (**Figure 2B, controls)**. Two independent stocks of the N-terminus truncated protein, HDM20-1 and HDM20-2, have 95% defective mitoses similar to the *orc6* deletion **(Figure 2B, N-terminal deletion)**. Stocks expressing Orc6 with a deletion of the middle portion of the protein, HDdel-1 and HDdel-2, also exhibit extensive cytological defects **(Figure 2B, Exon skipping)**. Transgenic flies expressing Orc6 with a wild-type human intron 4 show moderate improvement, with 50% (HDIn4-1) and 30% (HDIn4-2) of mitoses having normal configuration (**Figure 2B, Intron4 wild type)**. However, splice site mutation c.449+5G>A (HDIn4G>A-1 and HDIn4G>A-2) completely abolishes this positive dynamic **(Figure 2B, Intron4 mutant G>A)**.

**Figure 2.**
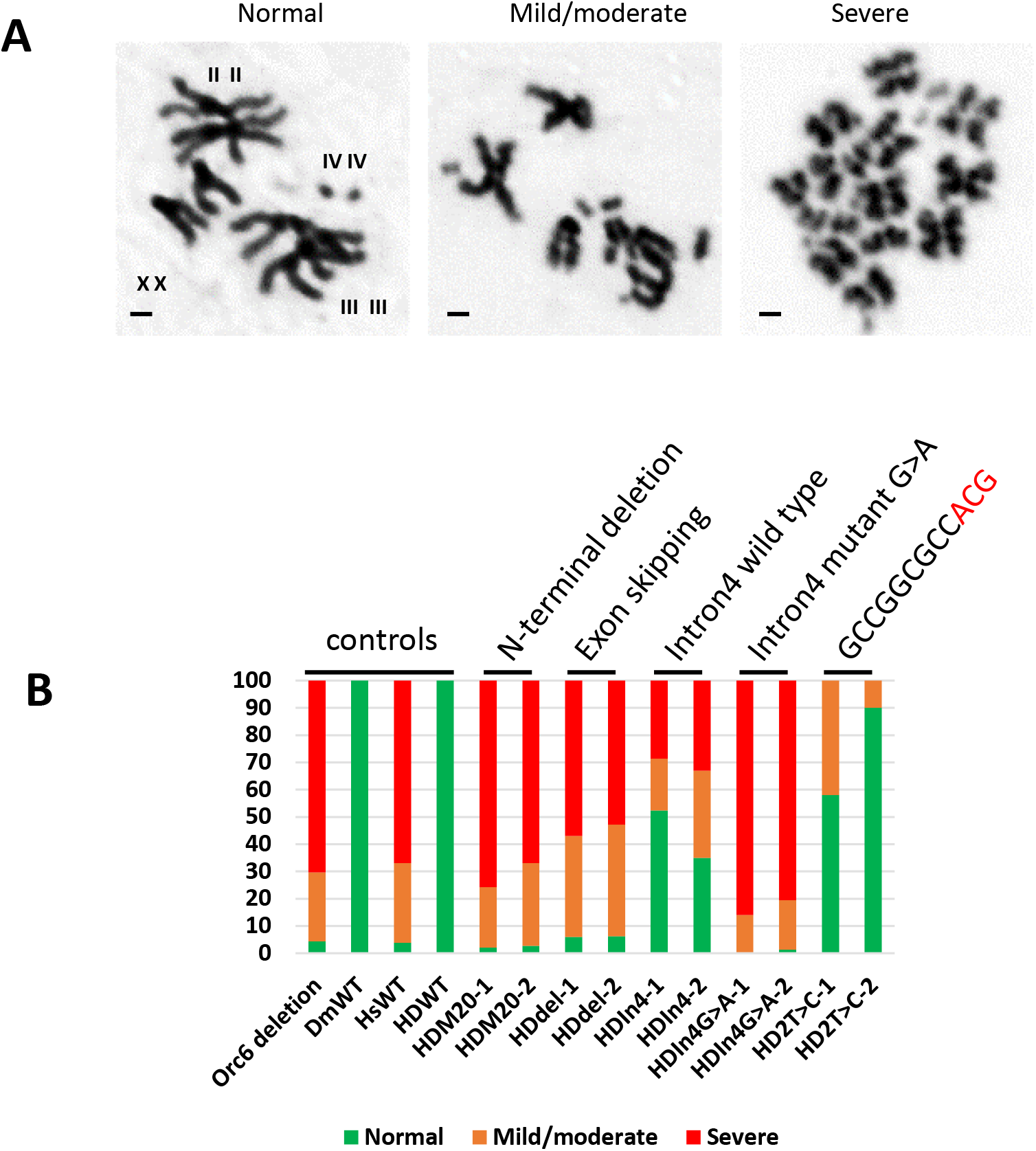
Analysis of chromosomal defects in MGS human-*Drosophila* mutants. **(A)** The analyzed karyotypes were divided into three classes: normal – all chromosomes are clearly visible, no defects; mild/moderate - karyotypes are defective, some chromosomes can be identified, but contain rearrangements (deletions, duplications, bridges); severe - chromosomes are abnormally condensed and fragmented. “XX” - pair of the X, “II II” – second, “III III”-third, and “IV IV” - fourth chromosomes. Scale bar 1μm. **(B)** Percentages of normal (green) and defective (orange and red) karyotypes was plotted on a bar chart. Two independent transgenic fly stocks were studied for each type of mutant MGS construct. DmWT and HDWT represent positive controls. Orc6 deletion and HsWT represent negative controls.

While karyotype analysis provides convenient tool for evaluating chromosome integrity, it is not sufficient to assess progress in development. In our previous study, we showed that the elevated expression of Orc6 Y225S or Orc6 K23E transgene can rescue the *orc6* deletion phenotype ^18,20^. We investigated whether it would be the case for the mutated MGS Human-*Drosophila* hybrids described above. The UAS promoter present in all constructs allows increased *Orc6* transgene expression by crossing to flies bearing the *tubulin* promoter-driven GAL4 (see Materials and Methods). As summarized in **Table 1**, the expression of HDWT Orc6 transgene rescued *orc6* deleted flies, but the expression of all deletion mutants, HDM20-1, HDM20-2, HDdel-1 and HDdel-2, under strong *tubulin* promoter did not rescue flies to the adult stage. Importantly, the expression of all transgenic proteins was successfully induced in flies as shown by a Western blot **(Figure 1C)**. We concluded that the deletion of the N-terminal 19 amino acids or the deletion of the middle part of the protein resulted in completely inactivated Orc6. This phenotype was inconsistent with the relatively mild phenotypes of MGS patients carrying these mutations and warranted further investigation.

**Table1.** Rescue of the *orc6* deletion with different hybrid transgenes under control of *tubulin* promoter.

| Transgenic construct | Molecular modification associated with MGS syndrome | Rescue of the *orc6* deletion by the expression of transgene under *tubulin* promoter |
| --- | --- | --- |
| HDWT | Wild type (no modifications) | + 79% (420) |
| HDM20-1<br>HDM20-2 | Deletion of the first 19 amino acids (c.2T>C) | - 0% (245)<br>- 0% (415) |
| HDdel-1<br>HDdel-2 | Deletion of the middle domain 30 amino acids | - 0% (397)<br>- 0% (490) |
| HDIn4-1<br>HDIn4-2 | Wild type intron4 | + 91% (659)<br>+ 27% (896) |
| HDIn4G>A-1<br>HDIn4G>A-2 | Mutation in the intron4 (c.449+5G>A) | - 0% (881)<br>- 0% (901) |
| HD2T>C-1<br>HD2T>C-2 | GCCGGCGCC precedes start codon ACG | + 90% (803)<br>+ 62% (453) |

Percentages of rescued flies were calculated based on expected segregation of phenotypes. Total progeny analyzed is shown in parentheses. (+), the expression of the transgene restores viability of *orc6*-deleted flies to adult stage. (-), no rescue.

Using the same rescue approach, we explored the possibility of ORC6 MGS mutation in intron 4, c.449+5G>A, to reduce splicing efficiency. Two independent transgenic fly stocks HDIn4-1 and HDIn4-2 carrying Human-*Drosophila orc6* gene with the wild-type intron 4 rescued *orc6* deletion to viability when expressed under *tubulin* promoter (**Table 1**). The resulting adult flies were phenotypically normal and undistinguishable from wild type flies as well as from HDWT. However, none of the mutant stocks HDIn4G>A-1 or HDIn4G>A-2 were able to rescue flies to adult stage (**Table 1**). The hybrid Orc6 protein was detected in flies carrying transgenes with wild type intron 4 (HDIn4-1 and HDIn4-2) but not intron 4 with mutation (HDIn4G>A-1 and HDIn4G>A-2) (**Figure 1D**). Since the sensitivity of the antibodies might be limited, in addition we analyzed RNA expression levels. We isolated total RNA from flies overexpressing hybrid Orc6 under *tubulin* promoter and performed RT-PCR reaction. The resulting cDNA was subjected to PCR analysis with primers specifically detecting only Orc6 HD hybrid. PCR product of the expected size was observed only for the transgene carrying the wild-type intron 4, indicating that the splicing of the mRNA was successfully completed **(Supplementary Figure 1A, HDIn4-1 and HDIn4-2)**. No PCR product was detected for transgene with mutant intron 4 (**Supplementary Figure 1A, HDIn4G>A-1 and HDIn4G>A-2)**. Since neither mRNA nor Orc6 protein was detected in the transgenes carrying the c.449+5G>A mutation, we concluded that this mutation completely abolishes splicing and therefore cannot be accountable for the observed phenotypes in MGS patients.

### The analysis of the DNA sequence context of the c.2T>C mutation

Our results indicate that both mutations under investigation, c.2T>C and c.449+5G>A, do not produce a functional Orc6 protein, which is inconsistent with the clinical phenotype reported for c.2T>C/c.449+5G>A case of ORC6 based MGS ^3^. To explore this paradox, we analyzed human *ORC6* mRNA sequence (https://www.ncbi.nlm.nih.gov/nuccore/AF139658.1) and found that the start codon follows the strong translation initiation sequence GCCGGCGCC. In 1989 Marilyn Kozak demonstrated using *in vitro* system that the initiation is very efficient when GCCGCCGCC precedes the AUG codon ^23^. We hypothesized that the strong upstream Kozak sequence facilitates the initiation of translation from the mutated ACG (c.2T>C) codon leading to the production of a full length Orc6 protein that could explain mild phenotype in MGS patients. To test this possibility, we mimicked the patient’s mutation DNA context by placing the GCCGGCGCC Kozak sequence upstream of the mutated c.2T>C start codon in the human*-Drosophila* Orc6 hybrid **(Figure 3A)**. The obtained two independent transgenic fly stocks rescued *orc6* deletion when expressed from native or *tubulin* promoter (**Figure 3A, Table 1**). As expected, the hybrid Orc6 protein was detected in rescued flies expressing protein under either native *orc6* or inducible *tubulin* promoters (**Figure 3B**). In our previous studies we found that flies carrying Orc6 based MGS mutations have missing or defective scutellar bristles ^18,20^. This specific phenotype was also observed in flies rescued with the transgene carrying GCCGGCGCC<u>ACG</u>G Kozak sequence (**Figure 3C, D, D’**), indicating that mutation c.2T>C is non-lethal and creates phenotype similar to other MGS mutations analyzed in flies.

**Figure 3.**
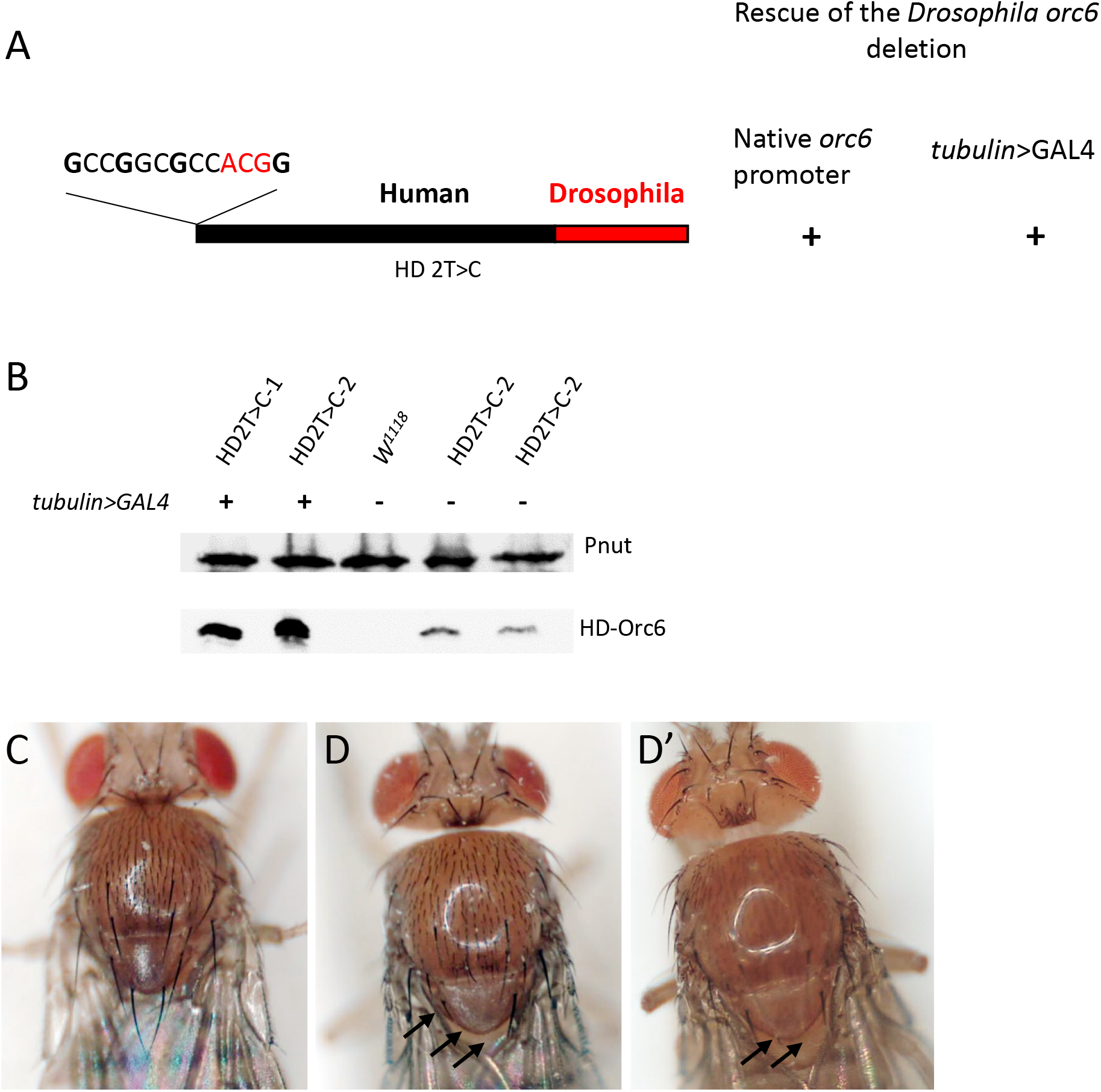
Mimicking the DNA sequence context of the Meier-Gorlin mutation c.2T>C in the Human-*Drosophila* hybrid. **(A)** Mutant start codon ACG embedded in strong Kozak sequence. Purines in positions +4, -3, -6, -9 are shown in bold. **(B)** Western blot of the flies carrying hybrid human-*Drosophila* transgenes. Two independent transgenic fly stocks, HD2T>C-1 and HD2T>C-2, were analyzed. (+) GAL4 boosts expression under *tubulin* promoter, (−) expression under native *orc6* promoter. *w*^*1118*^ – negative control (wild type flies). Pnut protein was used as a loading control. **(C**,**D**,**D’)** Phenotypes of *orc6* deletion flies rescued with HD2T>C transgene. **(C)** HDWT, wild type. **(D, D’)** HD2T>C, GCCGGCGCC precedes start codon ACG. Missing or defective scutellar bristles are pointed by arrows.

### The effect of Kozak sequence on the expression of the Orc6 Human-*Drosophila* hybrid

Next, to verify the importance of the upstream Kozak sequence, we mutated conservative guanines of the Kozak sequence (**G**CC**G**GC**G**CC<u>ACG</u>**G**) in positions -3, - 6, -9 to cytosines (**C**CC**C**GC**C**CC<u>ACG</u>**G**) to reduce the translation efficiency as reported in ^23^. The hybrid human-*Drosophila orc6* transgenes with native and mutated sequences were cloned into PMT/V5-HisA vector under control of metallothionein promoter. *Drosophila* S2 tissue culture cells were transfected, and transcription induced the next day with 0.5 mM CuSO_4_. The protein level of Orc6 HD hybrids was significantly higher in transgenes with native **G**CC**G**GC**G**CC<u>ACG</u>**G** sequence (**Figure 4A**). To confirm transgenes expression, we performed RT PCR of the total RNA from the transfected cells, and resulting cDNA was used in PCR reaction with primers specifically detecting Orc6 HD hybrids. The PCR product of expected 800 base pair length was detected in both native (**G**CC**G**GC**G**CC<u>ACG</u>**G)** and mutated (**C**CC**C**GC**C**CC<u>ACG</u>**G)** transgenes (**Figure 4B**) indicating that transcription efficiency was not affected. This result supports the idea that a strong Kozak context enhances the expression from the ACG codon.

**Figure 4.**
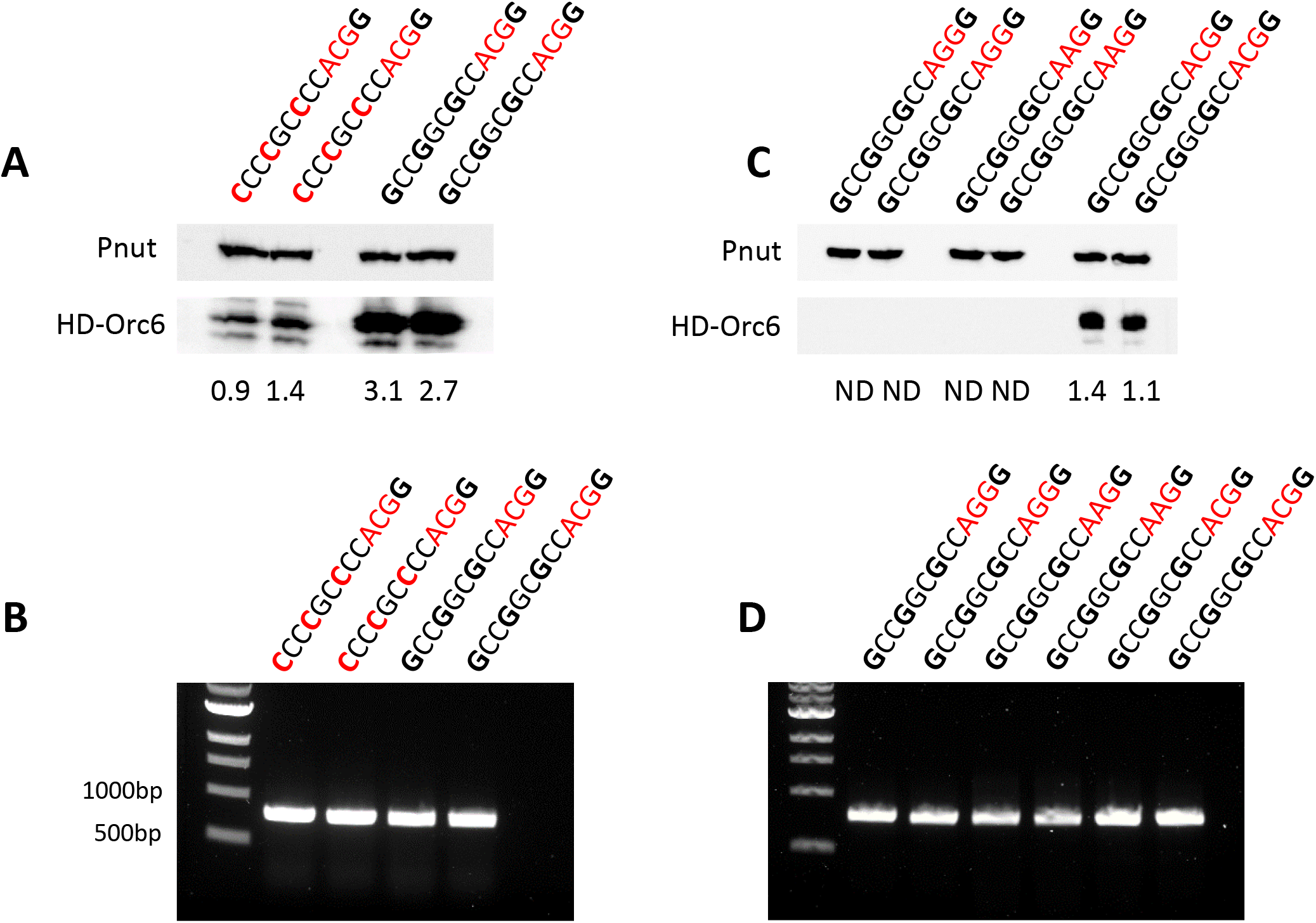
Expression of Human-*Drosophila* hybrid transgenes in S2 cells under metallothionein promoter. **(A)** Western blot of the two independent transfections with transgene carrying Kozak sequence **G**CC**G**GC**G**CCACG**G** and two independent transfections with transgene carrying Kozak sequence **C**CC**C**GC**C**CCACG**G** with mutated purines in positions -3, -6, -9. Pnut protein was used as a loading control. **(B)** PCR reactions of the cDNA synthesized from the transfected S2 cells mentioned above with primers detecting HD-Orc6 hybrids. **(C)** Western blot of the two independent transfections for each type of non-canonical codon: ACG, AAG, AGG. Pnut protein was used as a loading control. The numbers represent HD-Orc6: Pnut ratio, ND-not detected. **(D)** PCR reaction of the cDNA synthesized from S2 cells transfected with transgene carrying non-canonical codons.

While ACG has been shown to serve as a start codon with reduced efficiency compared to AUG, the AAG and AGG codons are not able to initiate translation ^27^. Therefore, if ACG works as initiator in our case, then the replacement of ACG with AAG or AGG will abolish the translation of the corresponding transgene. We mutated ACG to AAG or AGG in the same strong Kozak background: **G**CC**G**GC**G**CC<u>AAG</u>**G** and **G**CC**G**GC**G**CC<u>AGG</u>**G**. As shown in **Figure 4**, we could not detect any hybrid Orc6 protein from these constructs after transfection of the S2 cells (**Figure 4C**) even though the corresponding mRNAs were present (**Figure 4D**). This strongly supports the hypothesis that the non-canonical start codon ACG can initiate translation.

In summary, we conclude that in MGS patients with the c.2T>C/c.449+5G>A mutations in ORC6, the translation is initiated from a non-canonical ACG codon. The translation is governed by the strong Kozak sequence, which enables the recognition of ACG as an initiator codon and allows the production of the functional protein, albeit at a lower level, contributing to the observed MGS phenotypes.

## Discussion

Meier Gorlin Syndrome is a rare human disease associated with microcephaly, short stature, and multiple developmental defects ^2-4,28^. Mutations in a number of factors involved in DNA replication have been found to be causative for this disorder ^2,29-34^. According to a recent review, mutations in 13 DNA replication genes have been associated with MGS ^6^. All these genes have essential roles in DNA replication, and mutation in any of them could be potentially lethal. Therefore, elucidation of the molecular causes and clinical presentations for each gene is important for understanding pathology associated with these naturally occurring mutations.

Six mutations causing MGS are found in ORC6 ^6^. *Drosophila* and human ORC6 proteins have only 28% of sequence identity but are structurally similar and critical for the initiation of DNA replication ^14^. We have shown that mutations in conservative amino acids can be directly tested in *Drosophila*. In our previous works we successfully modeled Y232S (*Drosophila* Y225S), K23E and K202Rfs*1 (c.602_605delAGAA) MGS variants ^13,18,20^. The Y232S mutation impairs Origin Recognition Complex formation by reducing Orc6 interaction with subunit Orc3 ^16^. The K23E impairs DNA binding ^20^. The severity of abnormal embryological development in human K202Rfs*1 ^35^ is in agreement with a lethal embryonic phenotype in *Drosophila* for Orc6 having only the first 200 amino acids ^13^. During our molecular and genetic analyses of Orc6 functions we created human-*Drosophila* hybrid Orc6 protein (HD) containing intact human N-terminal domain (∼80% of the protein length) and *Drosophila* C terminus, which rescued Orc6-deficient flies to viable adults phenotypically undistinguishable from wild-type animals ^20^. This hybrid gene allows introducing mutations unique for human protein directly into human part of the hybrid and investigating the effect of mutations in live animal models. There are 92 *ORC6* variants of uncertain significance found in the public archive ClinVar, (www.ncbi.nlm.nih.gov/clinvar). Many of these variants could be evaluated *in vivo* using the hybrid transgene approach.

In the current study we focused on the compound heterozygosity in the *ORC6* gene: c.2T>C (p.Met1Thr) and c.449+5G>A found in MGS patients. Both mutations can theoretically generate truncated Orc6 protein variants with residual functionality ^6^. Using human-*Drosophila* hybrid Orc6 we found that truncated isoforms are not functional. Instead, strong Kozak sequence facilitates the translation of a full-length ORC6 protein from a non-AUG start codon ACG (c.2T>C mutation). It has been demonstrated that ACG codon placed in a strong Kozak sequence background, is less efficient in translation initiation compared to AUG (1-10% depending on the system analyzed) ^28,40^. Therefore, clinical phenotype observed in MGS patients could result from the insufficient amount of ORC6 protein. To test this, we enhanced the expression of the c.2T>C mutant HD transgene and restored the MGS phenotype observed in mutant flies to the wild type animals (Table1). These results indicate that increasing the amount of mRNA available for translation initiation can effectively compensate for the inefficient initiation from the ACG codon. This finding may have important implications for the development of therapeutic strategies. Since non-AUG translation is known to be regulated differently than the canonical AUG initiation, pharmacological modulation of noncanonical initiation codons may offer a novel therapeutic approach. Takacs et al. screened over 55,000 natural compounds in yeast to assess their effect on a luciferase reporter and identified two compounds that enhanced initiation from the ACG start codon ^36^. A similar screening could be conducted in *Drosophila* using HD hybrid as a reporter. Enhancing translation from ACG codon could be beneficial for patients with the c.2T>C mutation.

An ACG codon is naturally used by viruses, and the efficiency of translation depends on the optimal Kozak sequence ^37^. In humans, the TRPV6 gene (TRP channel, vanilloid type 6) starts from the non-canonical ACG triplet placed in a good Kozak context ^38^. Therefore, it is not surprising that Kozak sequence at the start of *ORC6* gene facilitates translation from ACG codon and compensates for the otherwise deleterious effect of the c.2T>C mutation. Recent bioinformatics analysis of thousands of human non-AUG extended proteoforms detectable with Ribo-seq reveals only 30 already annotated human genes and 390 potential candidates ^39,40^. What is the reason for such huge overproduction? Does it mean that translation initiation is error prone and thousands of the non-AUG proteoforms are not functional? These questions await answers. In conclusion, our work presented here demonstrates that despite evolutionary differences fruit flies can be used successfully to decipher molecular mechanisms associated with human diseases.

---

**Supplementary figure 1.**
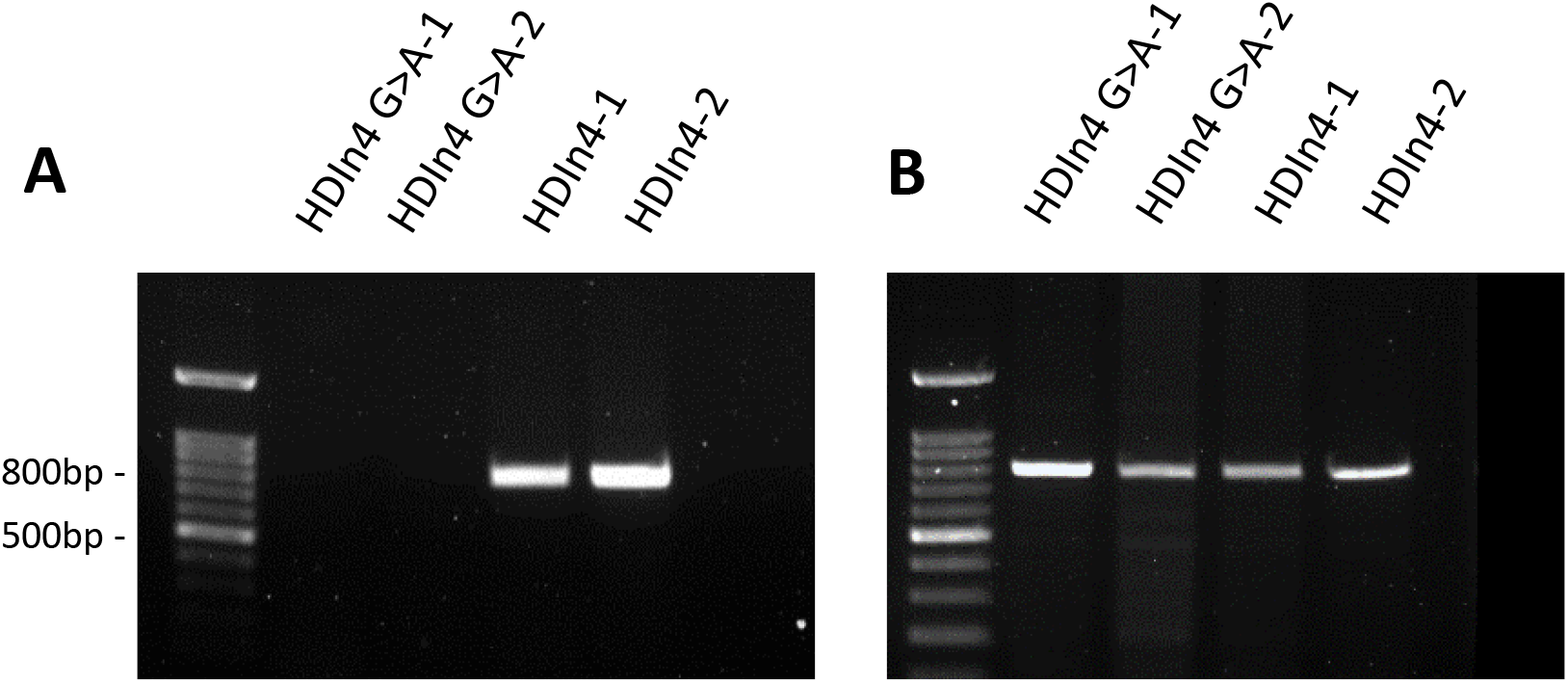
Expression of Human-*Drosophila* hybrid transgenes with intron 4 under control of the *tubulin* promoter. **(A)** PCR reactions of the cDNA synthesized from the polyA RNA isolated from two independent fly stocks bearing transgene with wild type intron 4 (HDIn4-1 and HDIn4-2) and two independent fly stocks with mutation in splice site c.449+5G>A (HDIn4 G>A-1 and HDIn4 G>A-2) with primers detecting HD Orc6 hybrid. **(B)** Control of the RT-PCR reactions. PCR reaction of the cDNA from the same samples was ran with primers detecting endogenous *Drosophila* Orc6. Heterozygous *orc6*^*35*^*/Cy;HDIn4/tub-GAL4* or *orc6*^*35*^*/Cy;HDIn4G>A/tub-GAL4* were used for cDNA preparation.

